# Characterization of the placenta-specific EnvV2-Fca soluble protein derived from endogenous retrovirus

**DOI:** 10.1101/2024.01.28.577663

**Authors:** Didik Pramono, Kenji Sugimoto, Tohru Kimura, Ariko Miyake, Kazuo Nishigaki

**Affiliations:** Laboratory of Molecular Immunology and Infectious Disease, Joint Graduate School of Veterinary Medicine, Yamaguchi University, 1677-1 Yoshida, Yamaguchi 753-8515, Japan; The Joint Graduate School of Veterinary Medicine, Yamaguchi University, 1677-1 Yoshida, Yamaguchi 753-8515, Japan

**Keywords:** endogenous retrovirus, retroviral *env*, placenta, domestic cat, evolution

## Abstract

Endogenous retroviruses (ERVs) are remnants of ancestral viruses in the host genome. Here, we identified the expression of a defective retroviral *env* gene belonging to the ERV group V member 2 Env (EnvV2) in *Felis catus* (EnvV2-Fca), which was specifically detected in the placental trophoblast syncytiotrophobic layer. EnV2-Fca was expressed as a secreted protein in cultured cells. Genetic analyses indicated that EnvV2 genes are widely present in vertebrate, and are under purifying selection among carnivores, suggesting a potential benefit for the host. Notably, this study suggests that birds, bats, and rodents carrying EnvV2 may play significant roles as intermediate vectors in spreading or cross-transmitting viruses among species. Overall, our findings provide valuable insights into the evolution of ERV in vertebrate hosts.

## Introduction

Endogenous retroviruses (ERVs) are ancient retroviral sequences that infect germ cells and are vertically transmitted to offspring in a wide range of vertebrate hosts, including mammals (1–4). Over time, this process generates a substantial proportion of ERVs in the mammalian genome via repeated germline reinfection (5). ERVs are inactivated through methylation and the accumulation of mutations, including nucleotide substitutions, deletions, and insertions (6). Over time, changes in viral genes can affect the function of the encoded proteins (7, 8). The proviral genome consists of three major polyproteins: *gag-pol*, and *env* (9). The *env* genes encode surface (SU) and transmembrane (TM) subunits that facilitate targeting and entry into specific cell types for infection (10). Previous research has demonstrated that defective *env*-ERV may play a critical role in physiological functions in the host, such as maintaining homeostasis, placentation, and acting as a restricting factor for infection (11–13). Defective *env*-ERV refers to an Env protein that has a signal peptide and SU subunit but contains a premature stop codon and lacks or has a partial deletion of the TM subunit.

It has been recently considered that defective *env*-ERV may function as an antiviral restriction factor (14, 15). In domestic cats, Refrex-1, derived from the defective *env* genes of ERV-DC7 and ERV-DC16, protects against feline leukemia virus (FeLV) subgroup D (FeLV-D) and ERV-DC Genotype I infections (14). In addition to its restrictive function, Refrex-1 maintains cellular homeostasis by regulating copper levels (11), and is fixed in the domestic cat population (14). Suppressyn is found in humans and is specific to the placenta (15, 16). Suppressyn, an *env*-ERV derived from human endogenous retrovirus (HERV)-F, contributes to the regulation of Syncytin-1 (the full-length *env* gene) involved in placentation (13), and has recently been demonstrated to have antiviral activity that restricts infection by mammalian type D retroviruses (15). In principle, defective Env-ERV protein expression can restrict viral infection by competing with exogenous Env for entry into host cell receptors during viral entry. This interaction between the defective Env-ERV protein and host receptor can protect host cells from viral Env protein invasion by controlling viral entry (10, 12, 15). These reports highlight the important function of defective *env*-ERV in hosts.

Other ERVs have been identified and expressed in mammals as placenta-specific proteins. The most well-known is syncytin, a captive retroviral envelope protein involved in human placental function (17). Along with syncytin, another placenta-specific protein, known as group V member ERV Env (EnvV), has been identified. EnvV can be classified into two groups, EnvV1 and EnvV2, which share high similarity (18). Initially, EnvV in humans (EnvV-Hum) was identified and neither EnvV1 nor EnvV2 exhibited fusion activity. The open reading frame (ORF) of EnvV1 comprises 477 amino acids, while the ORF of EnvV2 contains 535 amino acids (19). A single nucleotide insertion causes a truncation event at the C-terminus of genes, resulting in a frameshift (18). As previously reported, EnvV1- and EnvV2-encoded proteins are highly conserved during primate evolution (20). EnvV2 is intact in all analyzed primate species, whereas EnvV1 is preserved only in chimpanzees, gorillas, gibbons, rhesus macaques, baboons, African green monkeys (AGMs), tamarins, and saki monkeys. In these species, EnvV1 exhibits a full-length ORF with no C-terminal truncation or stop codons. The EnvV2 ORF in primates has been conserved since its integration over 40-45 million years ago and provides evidence of purifying selection of the EnvV2 gene among primates. Additionally, EnvV2 in macaques (EnvV2-Mac) exhibits fusion activity, but not with EnvV1 (20). Recently, after performing a genetic analysis using publicly available data, Simpson et al. (21) reported that EnvV1 and EnvV2 are highly conserved in two mammalian orders, Artiodactyla and Carnivora, suggesting that cross-species transmission occurred between these two orders at least 60 million years ago.

In this study, we focused on identifying defective *env*-ERV genes that may be beneficial to the host. We performed RNA-seq analysis using feline ovarian tissue and successfully detected the expression of retroviral elements. This element was identified as EnvV2 *Felis catus* (EnvV2-Fca). This study provides a comprehensive characterization of EnvV2-Fca, detailing the identification of the EnvV2 gene, the investigation of its expression and possible function, and an evolutionary analysis of EnvV2 in mammalian and non-mammalian vertebrate hosts.

## Materials and Methods

### Animals and Sampling

The domestic cat sample used in this study has been previously described (14). Briefly, the Nippon Institute for Biological Science provided a 2-month-old female specific-pathogen-free (SPF) cat that was euthanized, and an autopsy was performed. Animal tissues were stored at −80 °C until DNA or RNA was extracted for further investigation. Placental tissue was kindly provided by Dr. Daigo Umehara (Roji animal clinic, Fukuoka, Japan).

### RNAseq analysis using ovarian tissue

Total RNA was extracted from the ovaries of an SPF cat using a previously described method (14). The RNA-seq analysis was performed at the Yamaguchi University Genetic Experiment Facility (Yamaguchi, Japan).

### Cell lines

Cells were cultured in Dulbecco’s modified Eagle’s medium (DMEM; FUJIFILM Wako Pure Chemical Corporation, Osaka, Japan) supplemented with 10% fetal calf serum (FCS) and 1X penicillin-streptomycin. The cells were incubated in a CO_2_ incubator at 37 °C. The following cell types were used in this study: HEK293T (human embryonic kidney transformed with SV40 large T antigen) (22), 3201 (feline lymphoma cells) (23), 3281 (feline lymphoma) (24), FL74 (feline lymphoma) (25), FT-1 (feline T-cell leukemia) (26), MS4 (feline B-cell lymphoma) (27), KO-1 (lymphoma cell) (27), feline mammary adenocarcinoma cells (FMC, FKNp, FON, FRM, FNNm) (28), CrFK (feline kidney cells) (29), Fcwf-4 (feline macrophage-like cells) (30), G355 (feline fetal brain) (31), Fc9 (feline fetus), and Fet-J (feline peripheral lymphocytes) (26).

### PCR

The KOD-One Polymerase Kit (Toyobo, Japan) was used for various cloning processes. Primers were designed based on unique sequences both outside and inside each provirus to preserve the target gene. The primers used in this study are listed in Table 1.

**Table 1.**
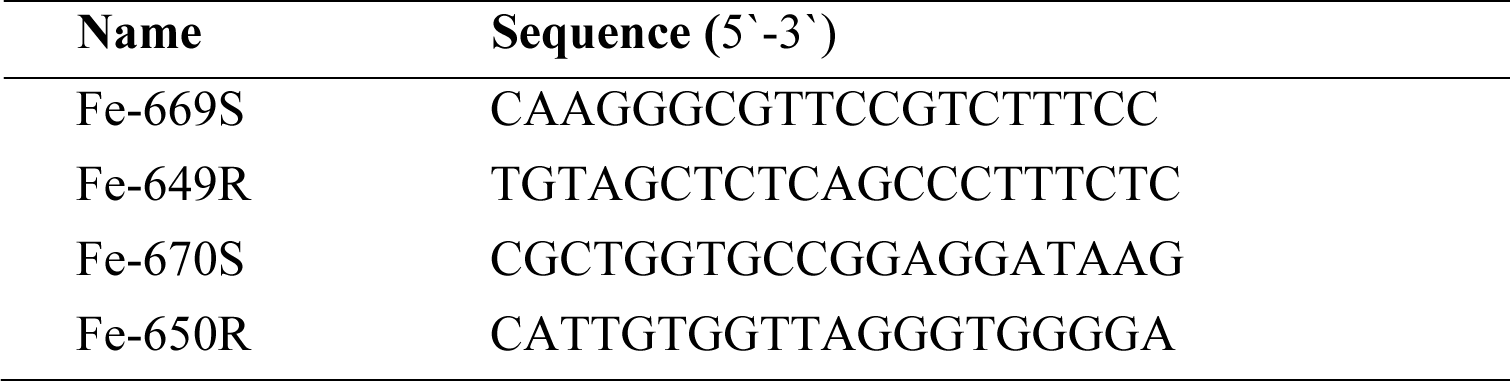
Primer information used for cloning EnvV2-Fca (domestic cat)

### Construction of expression vectors

The pFUΔss expression vector was used to construct the expression plasmids, as previously described (32). Each gene was PCR amplified from its respective plasmid using specific primers and enzyme sites. The resulting PCR products were subsequently digested with their corresponding restriction enzymes and cloned into the pFUΔss expression plasmid. The expression plasmid for EnvV2-Fca was constructed from domestic cat tissue cDNA using the primer pairs listed in Table 1. The plasmid was then inserted into the pFUΔss vector. The EnvV2-Mac sequence was obtained from the NCBI database and synthesized by Eurofins Genomics (Tokyo, Japan). The EnvV2-Mac gene was then inserted into the pFUSE-hIgG2-Fc2 vector (InvivoGen, California, USA). A myc tag was added to the C-terminus of EnvV2-Fca and EnvV2-Mac. The resulting expression plasmids were confirmed using sequencing (Fasmac Corporation, Atsugi, Japan).

### Transfection

HEK293T cells were transfected with expression plasmids using the TransIT-293 transfection reagent (Takara, Shiga, Japan), following the manufacturer’s instructions.

### Immunoblotting

Plasmids were introduced into HEK293T cells using the TransIT®-293 (Takara) reagent in six-well plates. After two days, the cell pellets were collected by washing three times with phosphate buffered saline (PBS). To prepare cell lysates, cells were mixed with lysis buffer containing 20 mM Tris-HCl (pH 7.5), 150 mM NaCl, 10% glycerol, 1% Triton X-100, 2 mM EDTA, 1 mM Na3VO4, and 1 µg/ml of aprotinin and leupeptin. The lysates were then placed on ice for 20 min. Insoluble components were removed by top-speed centrifugation (15,400 × g) for 20 min at 4 °C. The protein concentrations were calculated using a protein assay kit (Bio-Rad, Hercules, CA, USA).

The cell supernatants were filtered using a 0.22-µm filter and subsequently incubated with the desired antibody overnight at 4 °C. The purified protein, which included both the cell lysate and supernatant, was mixed with Sample Buffer Solution (Nacalai Tesque, Kyoto, Japan) and heated at 95–100 °C for 5 min. Sodium dodecyl-sulfate polyacrylamide gel electrophoresis (SDS-PAGE) was performed using 4–20% gels (Invitrogen, Carlsbad, CA, USA) at 100 V for 2 h. The gels were then transferred onto nitrocellulose membranes for western blot analysis. The following primary antibodies were used in the assays: anti-Myc monoclonal antibody conjugated with horseradish peroxidase (FUJIFILM Wako Pure Chemical Corporation; dilution, 1:1000). The substrate used was LumiGLO® Reagent (20×) and 20X peroxide (Cell Signaling Technology in Danvers, Massachusetts). Blots were imaged using Amersham ImageQuant 800 (Cytiva, Shinjuku, Japan).

### Detection of EnvV2-Fca expression via qRT-PCR

Total RNA was extracted from the tissues of a SPF cat (14), as well as from various feline cell lines using an RNAiso Plus kit (Takara) according to the manufacturer’s instructions. cDNA was synthesized using a PrimeScript II first-strand cDNA synthesis kit (Takara) following the manufacturer’s instructions. Prior to reverse transcription, the RNA samples were treated with recombinant DNase I (TaKaRa). The cDNA was amplified using Premix Ex Taq (Probe qPCR; Takara) in a CFX96 Touch Real-Time PCR Detection System (Bio-Rad, Hercules, CA, USA). The *EnvV2-Fca* gene was amplified using primers Fe-gvm2Env-F and Fe-gvm2Env-R and detected using probe Fe-gvm2Env-P (containing 6-carboxy-fluorescein; FAM) (Takara). The internal control, feline peptidyl prolyl isomerase A (*PPIA*), was amplified with primers Fe-227S and Fe-204R using SYBR Premix Ex Taq II (Tli RNaseH Plus; Takara) (Table 2). Thermal cycling was performed according to the manufacturer’s instructions.

**Table 2.**
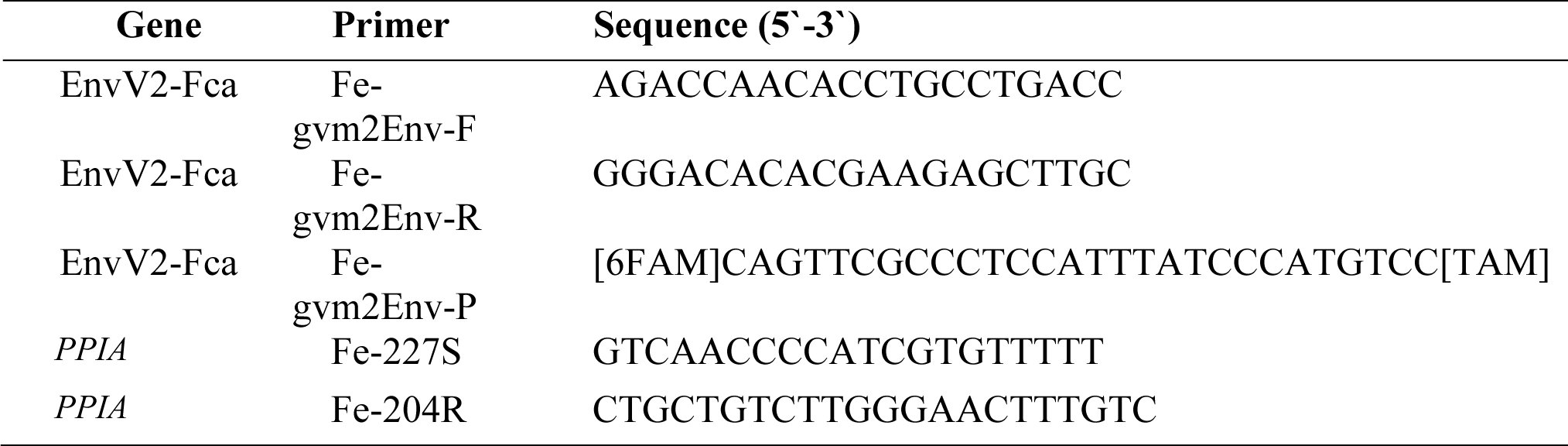
Primer and probe information used for qRT-PCR.

### In situ hybridization (ISH) analysis of placental sections

Fresh placental samples were fixed in situ, extracted, and dried onto microscope slides in preparation for RNA-ISH analysis. Fixed tissues were embedded in paraffin on CT-Pro20 (Genostaff, Tokyo, Japan) using G-Nox (Genostaff) as a less toxic organic solvent and sectioned into 6-µm thick sections. ISH analysis was performed with an ISH Reagent Kit (Genostaff) according to the manufacturer’s instructions. Tissue sections were deparaffinized with G-Nox and rehydrated in ethanol and PBS. The sections were then fixed with 10% neutral buffered formalin (NBF) for 30 min at 37 °C and washed in distilled water, placed in 0.2% HCl for 10 min at 37 °C and washed in PBS, treated with 4µg/ml ProteinaseK in PBS for 10 min at 37 °C and washed in PBS, and finally placed within a coplin jar containing 1x G-Wash (Genostaff) equal to 1x saline-sodium citrate (SSC). Hybridization was performed with probes (250 ng/mL) in G-Hybo-L (Genostaff) for 16 h at 60 °C. After hybridization, the sections were washed three times with 50% formamide in 2x G-Wash for 30 min at 50 °C and five times in TBST (0.1% Tween20 in Tris-buffered saline) at room temperature. After treatment with 1x G-Block (Genostaff) for 15 min at room temperature, the sections were incubated with anti-DIG AP conjugate (Roche; 1:2000) with G-block (diluted 1/50) in TBST for 1 h at room temperature. The sections were washed twice with TBST and incubated in 100 mM NaCl, 50 mM MgCl2, 0.1% Tween20, and 100 mM Tris-HCl (pH 9.5). Coloring reactions were performed with nitro blue tetrazolium/5-Bromo-4-chloro-3-indolyl phosphate (NBT/BCIP) solution (Sigma), and then sections were washed in PBS. Sections were counterstained with Kernechtrot Stain Solution (Muto), and mounted with G-Mount (Genostaff) and malinol (MUTO PURE CHEMICALS, Tokyo, Japan). Images were obtained using a NanoZoomer S210 digital slide Scanner: C13239-01 (Hamamatsu Photonics, Shizuoka, Japan) and NDP.view2 Plus Viewing Software: U12388-02 (Hamamatsu Photonics).

### Functional assay of EnvV2-Fca and EnvV2-Mac into cells

HEK293T cells were transfected with the expression vectors of EnvV2-Fca and EnvV2-Mac using TransIT®-293 Trasfection Reagent, following the manufacturer’s instructions. HEK293T cells transfected with a pFUΔss empty vector were used as a negative control. The cells were cultured for 2 days, and the culture supernatant was removed and stained with Diff-Quick staining solution according to the manufacturer’s instructions. Cell morphology was observed using an optical microscope to evaluate morphological changes.

### Phylogenetic and sequencing analysis

A phylogenetic tree was constructed using the sequences listed in the Accession Number section. The MEGA11 software package was used for phylogenetic analysis, and amino acid sequences were aligned using MUSCLE (33). A phylogenetic tree was constructed using the neighbor-joining method (34) and the amino acid substitutions JTT model (35), and robustness was evaluated through bootstrapping (1,000 times) (36). All packing was conducted using MEGA11 (37).

### Evolutionary Analysis of EnvV2 among carnivores

To conduct evolutionary analyses of EnvV2, we searched for publicly available genome data using the BLAST method. We searched the vertebrate genome database using the env ORF of EnvV2-Fca (carnivora), EnvV2-Mac (primate), and *Bos indicus* (even-toed ungulate) nucleotide sequences as references. We obtained EnvV2 sequences from carnivores and aligned them using MUSCLE (33). We then calculated the pairwise p-distance values of the amino acids, which represent the proportion of amino acid sites at which the two sequences differ. The number of non-synonymous substitutions (dN) and synonymous substitutions (dS) per site were estimated using the Nei-Gojobori method (38). The packing procedure was conducted using MEGA11 (37). To analyze amino acid similarity, the amino acids of EnvV2 Carnivores were aligned using MUSCLE, and the amino acid similarity (%) of all carnivore representatives was calculated using Genetyx 16 (Genetyx Corporation, Tokyo, Japan).

## Results

### EnvV2-Fca is a retroviral defected *env* gene in domestic cats

To identify a candidate for the retroviral *env* gene in domestic cats, RNA-seq analysis was performed using RNA obtained from the ovarian tissue of a domestic cat. We identified *env* gene fragments, classified as EnvV2, by focusing on genes with high expression levels. We isolated EnvV2-Fca from various domestic cat samples (spleen and small intestine) using RT-PCR. A positive band of approximately 2.2 kb was identified in the spleen and small intestine. We conducted molecular cloning of these PCR-positive samples to obtain the EnvV2-Fca ORF sequence. EnvV2-Fca has a nucleotide sequence of 1317 bp, encodes 439 amino acids, and contains a signal peptide (SP), surface unit (SU) subunit, furin cleavage site, fusion peptide, immunosuppressive domain (ISD), and truncated transmembrane (TM) subunit (Figure 1A). EnvV2-Fca is an Env protein that differs from retroviral envelope-derived proteins due to its truncated TM subunit structure. Furthermore, several mutations were observed compared to the reference data (predicted EnvV2-Fca) in the NCBI database (accession number XM_019831672). Figure 1B displays the changes in the amino acid positions. Phenylalanine at position 13 was replaced with leucine (F13L), methionine at position 41 with valine (M41V), threonine at position 139 with alanine (T139A), and glutamine at position 321 with arginine (Q321R).

**Figure 1.**
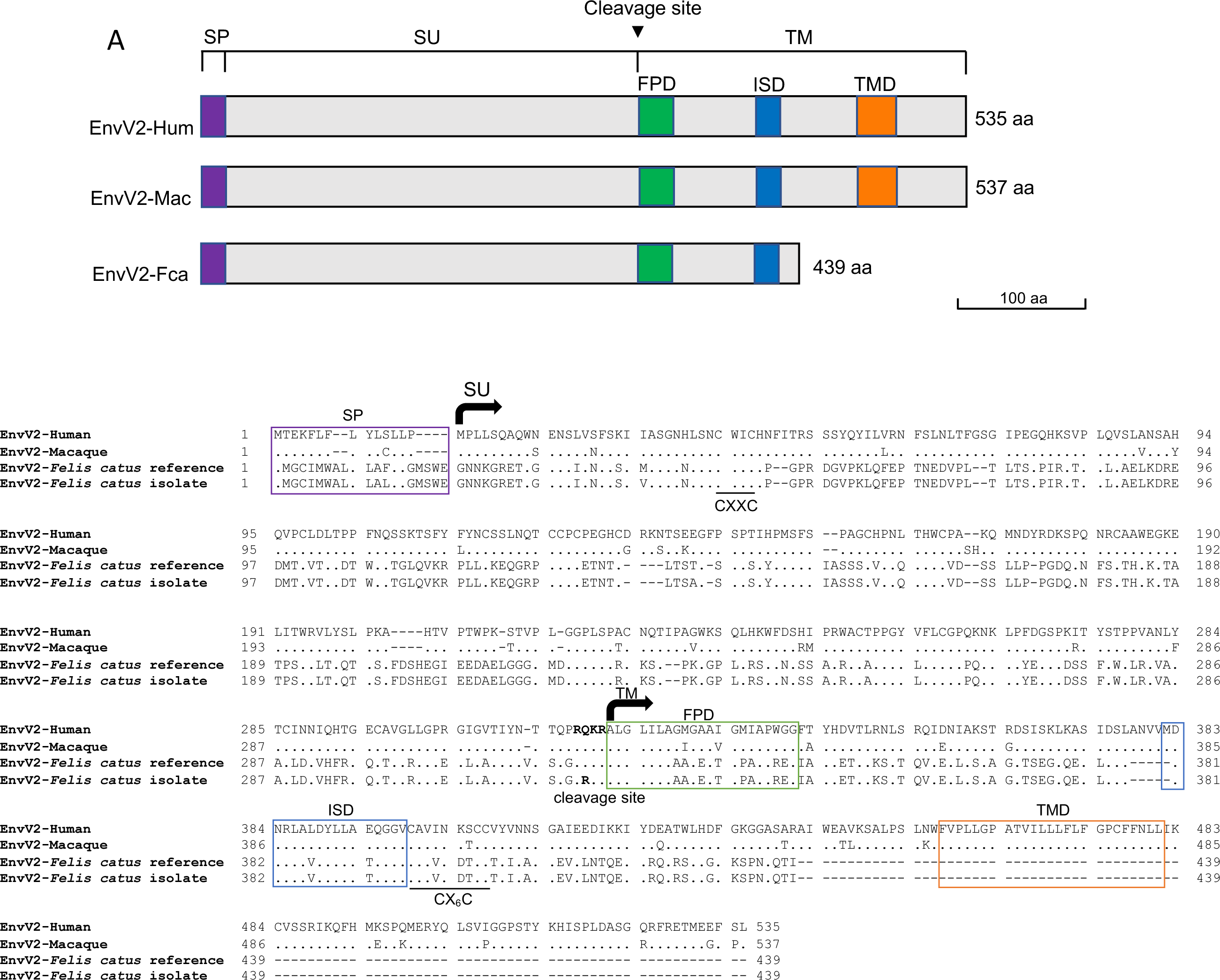
Structure schematic and amino acid alignment of EnvV2-Fca. (A) The schematic representation of the structure of EnvV2-Fca, EnvV2-Hum, and EnvV2-Mac. number of amino acids is indicated on the right side. (B) Amino acid sequence alignment of the Env proteins of EnvV2-Hum, EnvV2-Fca from NCBI (reference), and the isolate. SP, signal peptide; SU, surface unit; TM, transmembrane subunit; FPD, fusion peptide domain; ISD, immunosuppressive domain; TMD, trans-membrane domain; R-X-R/K-R is the cleavage motif; CXXC and CX_6_CC are sites of covalent interaction.

### EnvV2-Fca is expressed as a soluble protein

To determine whether the protein of the EnvV2-Fca gene can be expressed, we cloned the EnvV2-Fca ORF into the pFUΔss expression vector (32). A myc tag was added to the C-terminus of EnvV2-Fca. We then analyzed the expression of EnvV2-Fca by immunoblotting the supernatant and cell lysate from transfected HEK293T cells. Supernatants from transfected HEK293T cells containing EnvV2-Fca expression plasmids were immunoprecipitated using an anti-myc antibody and western blot analysis was performed. Figure 2A shows that EnvV2-Fca expression was detected in the cell lysates at a molecular mass of 55 kDa. However, the size of EnvV2-Fca in the cell supernatants was unexpectedly large, ranging from 60-70 kDa. This size difference may be due to various forms of protein modification. Based on the amino acid sequence, EnvV2-Fca was expected to contain a furin cleavage site (R-R-K-R) (Figure 1B); however, cleavage of the EnvV2-Fca protein was not observed when expressed in transfected HEK293T cells. Only a single band representing the protein expression of EnvV2-Fca was observed in both the cell lysates and supernatant (Figure 2A). These results indicate that EnvV2-Fca was secreted from the cells as a soluble protein, as these proteins were detected in the cell supernatants.

**Figure 2.**
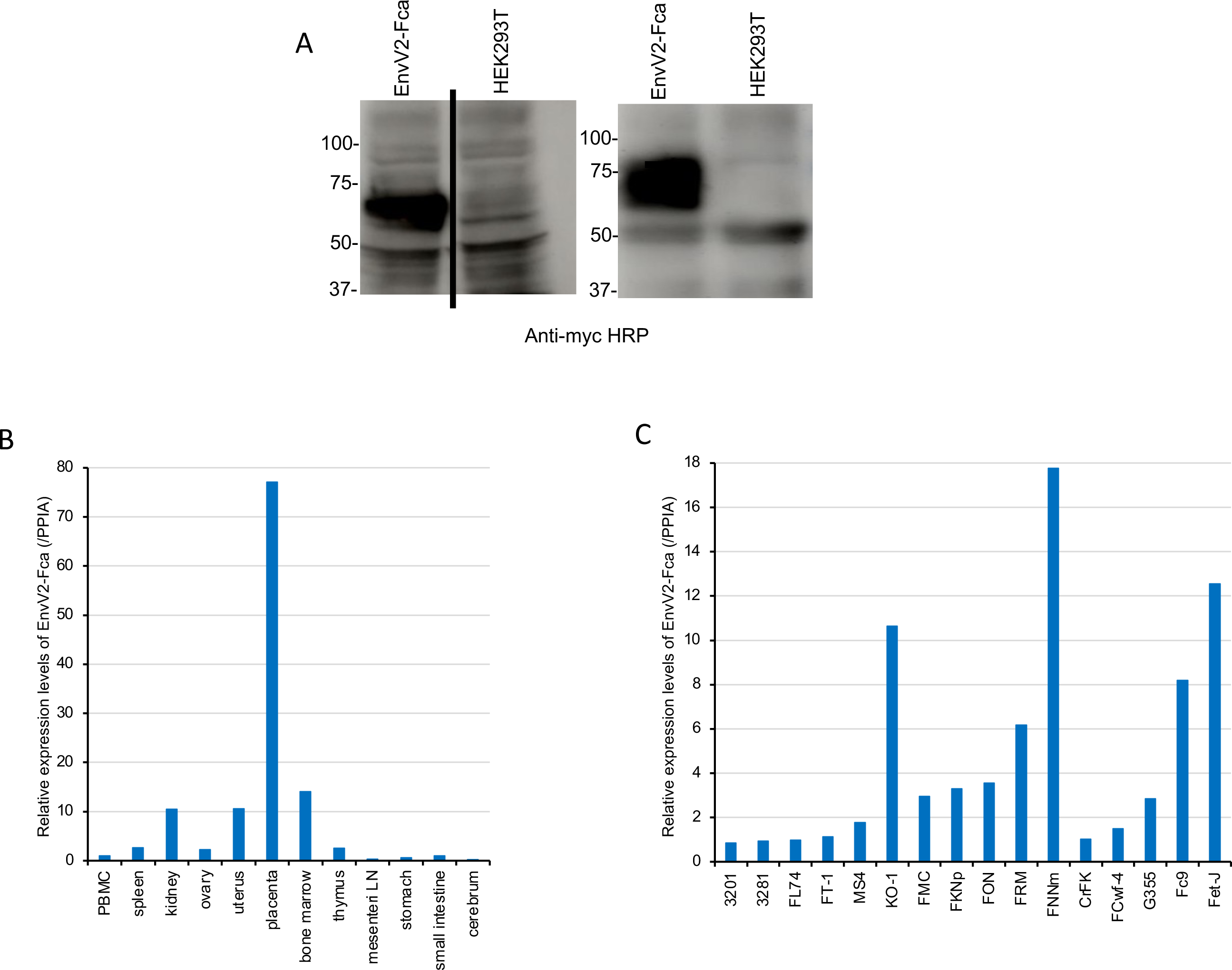
Expression of EnvV2-Fca. (A) The presence of the EnvV2-Fca protein was detected in both the cell lysates (left) and supernatants (right) from HEK293T cells transfected with the indicated plasmids. Immunoprecipitation (IP) and western blotting (WB) were used to analyze EnvV2 and the negative control using an anti-Myc antibody. (B, C) The expression of EnvV2-Fca was analyzed in feline tissues and cell lines. The EnvV2-Fca transcripts were quantified using quantitative RT-PCR in feline tissues and cell lines. The x-axis represents the analyzed samples. The expression level on the y-axis is normalized to the expression of peptidylprolyl isomerase A (*PPIA*). The normalized expression in feline tissues and cell lines is shown as 1 in peripheral blood mononuclear cells (PBMC). LN, Lymph node.

### Expression analysis of EnvV2-Fca in feline tissues and cell lines

To determine the tissue-specific expression of EnvV2-Fca, we conducted quantitative RT-PCR (qRT-PCR) to analyze the expression levels of EnvV2-Fca. We quantified the expression of EnvV2-Fca in normal feline tissues (specific-pathogen free cat) and cell lines. As shown in Figure 2B, EnvV2-Fca was broadly expressed in a variety of tissues, with the highest expression levels observed in the placental tissue. EnvV2-Fca was detected in all feline cell lines (Figure 2C). Our findings suggest that EnvV2-Fca is widely expressed in feline tissues and cell lines, with particularly high expression levels in the placenta. This suggests that EnvV2-Fca was expressed in a placenta-specific manner.

### In situ hybridization analysis (ISH analysis) of EnvV2-Fca in the placenta

To investigate the possible importance of specific EnvV2-Fca expression in the placenta, as shown by the qRT-PCR assay in Figure 2B, ISH was performed on paraffin-sectioned placental tissue. To detect EnvV2-Fca transcripts, specific digoxigenin-labeled antisense probes were developed, and sense probes were used as negative controls. As shown in Figure 3, specific labeling was observed only with antisense probes and not with the control probes. As shown in Figure 3 (right), the specific labeling of EnvV2-Fca was broadly distributed in the placenta, including the connective tissue, with a particularly significant signal in the trophoblast syncytiotrophobic layer. This finding indicates that EnvV2-Fca is expressed in the placental tissue and is specific to the placenta.

**Figure 3.**
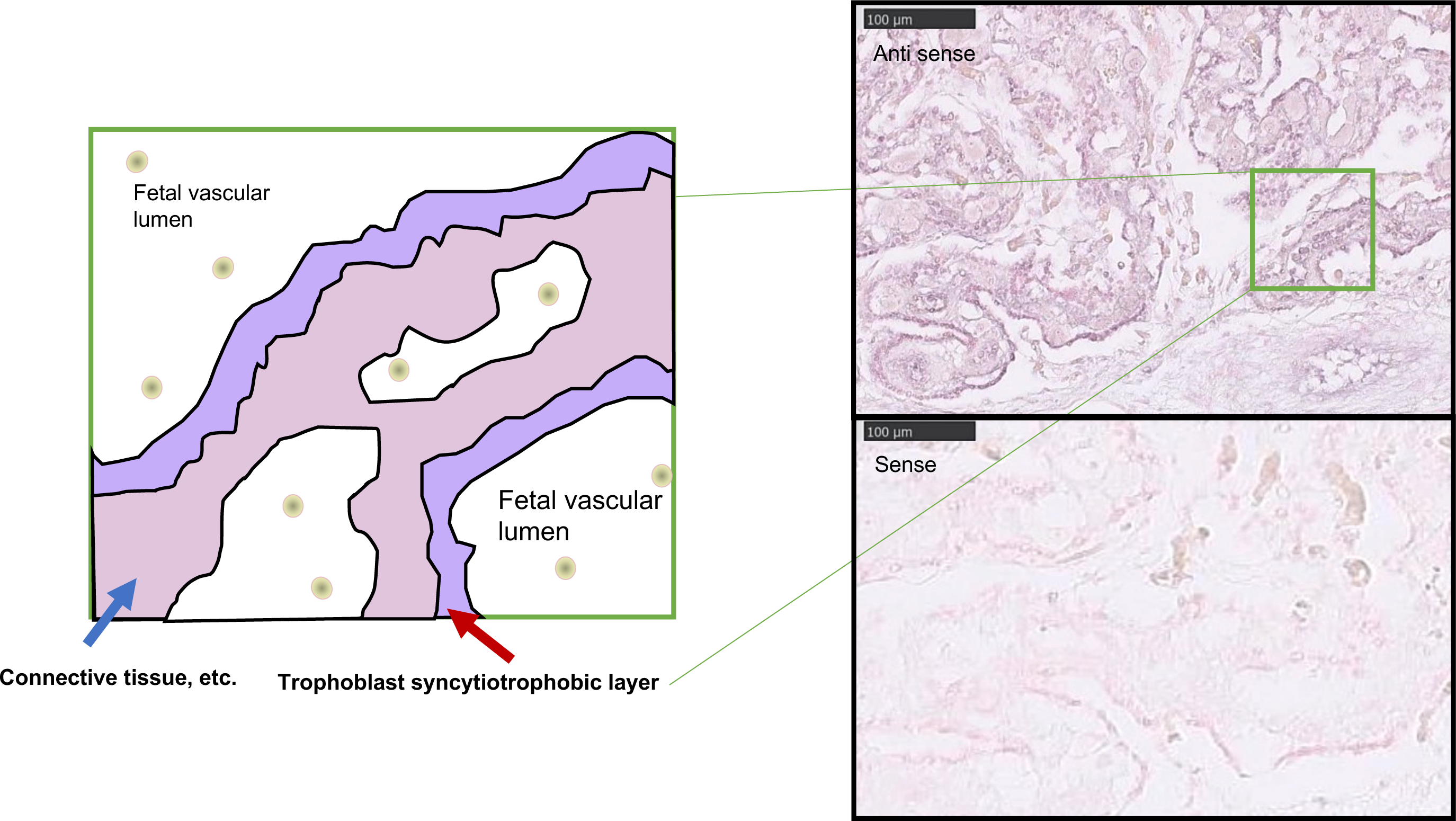
In situ hybridization analysis of EnvV2-Fca in the placenta. (A) Schematic representation of the cat placenta, which contains connective tissue, a fetal vascular lumen, and trophoblast syncytiotropobic layer. (B) In situ hybridization was performed on placental sections using a digoxigenin-labeled antisense (upper) or sense probe (negative control; lower). The labeled syncytiotrophoblast was observed at a higher magnification. (Scale bars=100 μm).

### EnvV2-Fca exhibits non-fusogenic activity

Previous studies have demonstrated that EnvV2-Mac has fusogenic activity and is predicted to play a fundamental role in syncytiotrophoblast cell-cell fusion (20). However, a comparison of the structures of EnvV2-Fca and EnvV2-Mac revealed that EnvV2-Fca has a truncated TM subunit (Figure 1A). Although EnvV2-Fca has a different structure, we observed that it is specifically expressed in the placenta (Figure 2 and 3). We subsequently investigated whether EnvV2-Fca has cell fusion activity, which is the hypothesized function of retroviral *env* genes that are specifically expressed in the placenta. A functional assay was performed by transfecting cell lines in culture with the abovementioned env-expression vectors, as previously described (39), and following up on syncytium formation two days post-transfection. The results indicated that EnvV2-Fca exhibited non-fusogenic activity compared to the positive control, whereas EnvV2-Mac exhibited fusion activity (20) (Figure 4). This suggests that despite EnvV2-Fca being expressed specifically in the placenta, it does not exhibit fusogenic activity.

**Figure 4.**
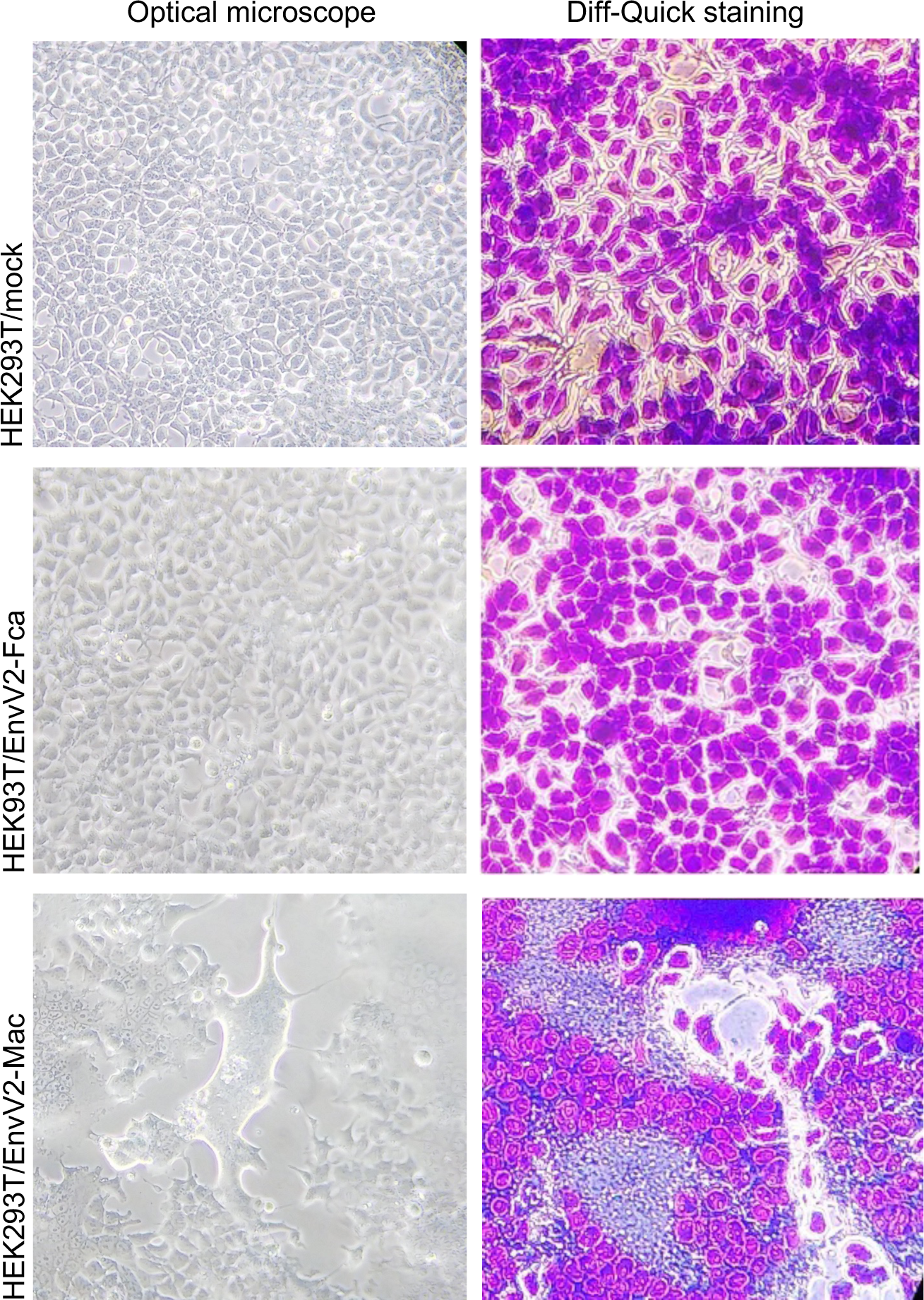
Functional assay of EnvV2-Fca and EnvV2-Mac. Morphology of HEK293T cells transfected with EnvV2-Fca, EnvV2-Mac, and mock (Empty vector). Images were obtained using optical microscopy, without staining (left), and after Diff-Quik staining (right) 24 h after transfection.

### Conservation and evolution of EnvV2 genes among mammals and non-mammalian vertebrates

The evolution of EnvV2 has been studied in several mammals, including humans, primates, carnivores, and even-toed ungulates (19–21). However, these reports also tend to mention EnvV1, which was not the focus of our study. We hypothesized that the evolution of EnvV2 is not limited to these mammals. We further investigated EnvV2 by analyzing publicly available sequence data using the *env* ORF of EnvV2-Fca (carnivora), EnvV2-Mac (primate), and *Bos indicus* (even-toed ungulate) as search references. In the NCBI database, we located an EnvV2 prediction labeled as ‘endogenous retrovirus group V member 2 Env polyprotein-like,’ which we refer to as EnvV2. We collected and analyzed all of the data labeled with EnvV2 from the NCBI database. Our analysis revealed that several additional mammals, including bats, rodents, and elephants, also contain EnvV2 (Figure 5A and 5B). We compared the amino acid similarity of these mammals to EnvV2-Fca and found that bats, rodents, and elephants share amino acid similarities of 60-68%, 47-71%, and 57% with EnvV2-Fca, respectively. To determine whether EnvV2 is conserved only in mammals, we conducted a search for publicly available data by performing a BLAST search on vertebrates. Our search revealed that EnvV2 also appears in several non-mammalian species, including birds, turtles, amphibian, fish, and reptiles (Figure 5).

**Figure 5.**
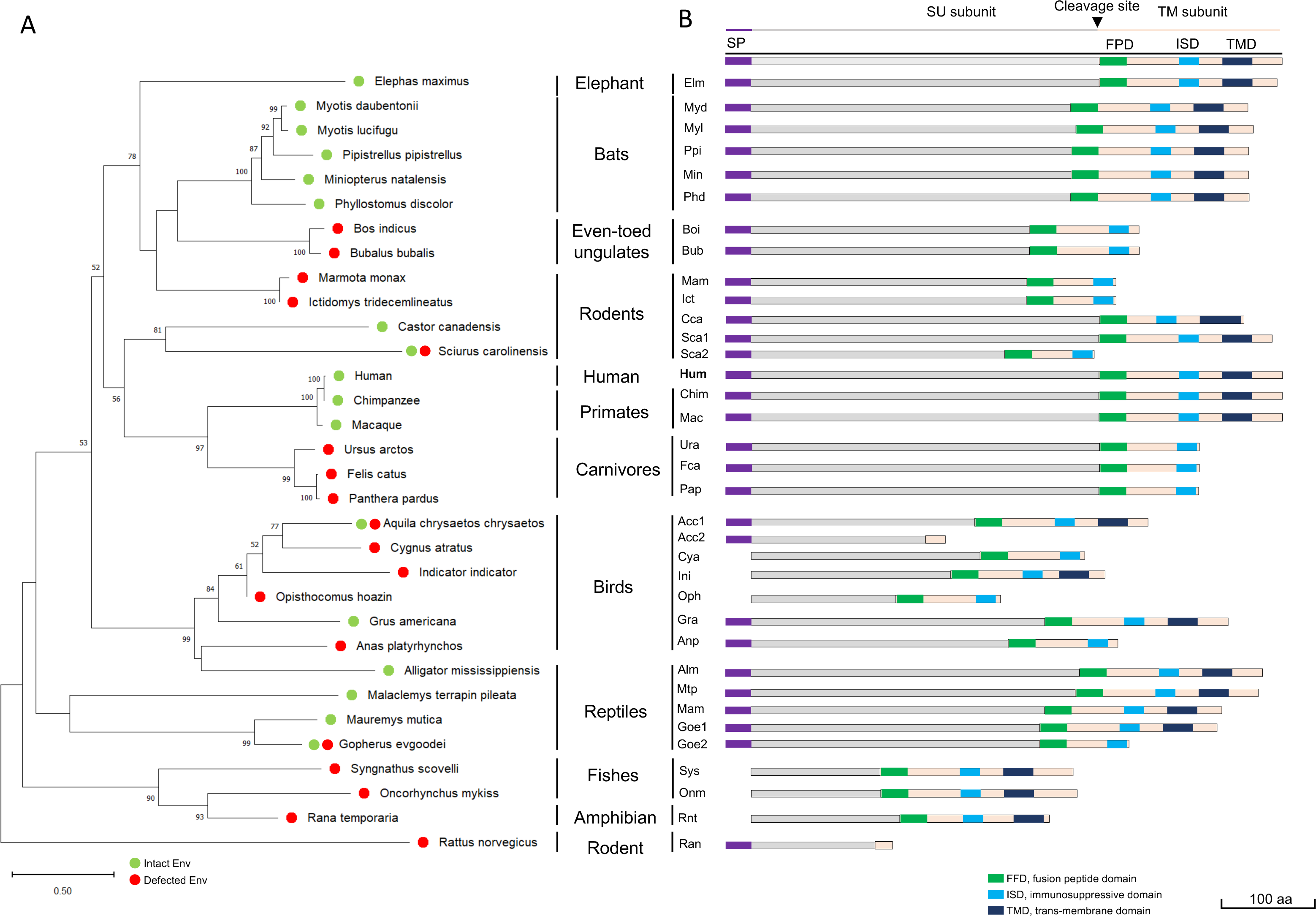
Phylogenetic tree and schematic of EnvV2 based on amino acids from mammalian and non-mammalian vertebrates. (A) The phylogenetic tree was constructed using the neighbor-joining method, based on EnvV2 amino acid sequences. The length of the horizontal branches is the percentage of amino acid substitutions from the node (left scale bar) and the percentage bootstrap values from 1,000 replicates are indicated at the nodes. (B) A schematic of EnvV2, which was classified based on amino acids from mammalian and non-mammalian vertebrates.

A phylogenetic tree was constructed using the amino acid sequence of EnvV2 ORF (Figure 5A). The tree revealed that the currently identified genes are distinct from previously identified EnvV2 genes, especially between mammalian and non-mammalian vertebrates, which are located separately. Notably, EnvV2 in bats, rodents, and ungulates belong to the same lineage (Figure 5A). Additionally, EnvV2-Fca, a carnivorous order, is located in the same clade as that of humans and primates. We conducted further analyses on the identified sequences. Our results revealed a characteristic structure of the retroviral protein (40–42) with a putative furin cleavage site, and a consensus of R/K-X-R/K-R representing the SU and TM subunits. However, several of the identified EnvV2 proteins had a modified furin cleavage site. Additionally, we observed a CX_2_C motif that corresponds to the binding domain between the two subunits and a CX_6_CC motif as an envelope glycoprotein predicted to facilitate fusion (42). Using Phobius (https://phobius.sbc.su.se/) (42), we identified a signal peptide at the N-terminal of the sequences. Based on sequence structure analysis, we classified EnvV2 complete Env (containing a signal peptide, SU, furin cleavage site, and TM subunits) as present in humans, primates, elephants, birds (*Aquila chrysaetos chrysaetos* and *Grus americana*), rodents *(Castor canadensis, Sciurus carolinensis*), and reptiles. However, we identified two defective EnvV2 in two groups: one EnvV2 with a truncated TM subunit present in Carnivores, even-toed ungulates, rodents (*Marmota monax, Ictidomys tridecemlineatus, Sciurus carolinensis,* and *Rattus norvegicus*), bird (*Aquila chrysaetos chrysaetos),* and reptiles (*Gopherus evgoodei*), and the other lacking a signal peptide or partial deletion of the SU subunit present in birds (*Cygnus atratus, Indicator indicator, and Opisthocomus hoazin*), fishes, and amphibia. Interestingly, all of the identified EnvV2 sequences conserved the immunosuppressive domain (ISD), with the exception of one rodent species (*Rattus norvegicus*).

Next, we analyzed the evolution of EnvV2. The evolution of EnvV2 can be investigated by comparing the rates of non-synonymous (dN) and synonymous (dS) changes in similar or related sequences (43). ERV genes, such as host genes, can be conserved in two ways: (a) when the dN/dS rate is less than 1, the genes are under purifying selection and retain a beneficial function. (b) If the dN/dS rate is greater than 1, the genes are classified under positive selection. The domestic cat (*Felis catus*), which belongs to the order Carnivora, was used to study the conservation and evolution of EnvV2-Fca in Carnivora species. We calculated the ratio of non-synonymous to synonymous mutations (dN/dS) using the method developed by Nei and Gojobori (38). The results suggested that the dN/dS ratios of EnvV2 among carnivores range from 0.08 to 0.81 (Figure 6). Our findings suggest that EnvV2 genes are subject to purifying selection in all Carnivora species.

**Figure 6.**
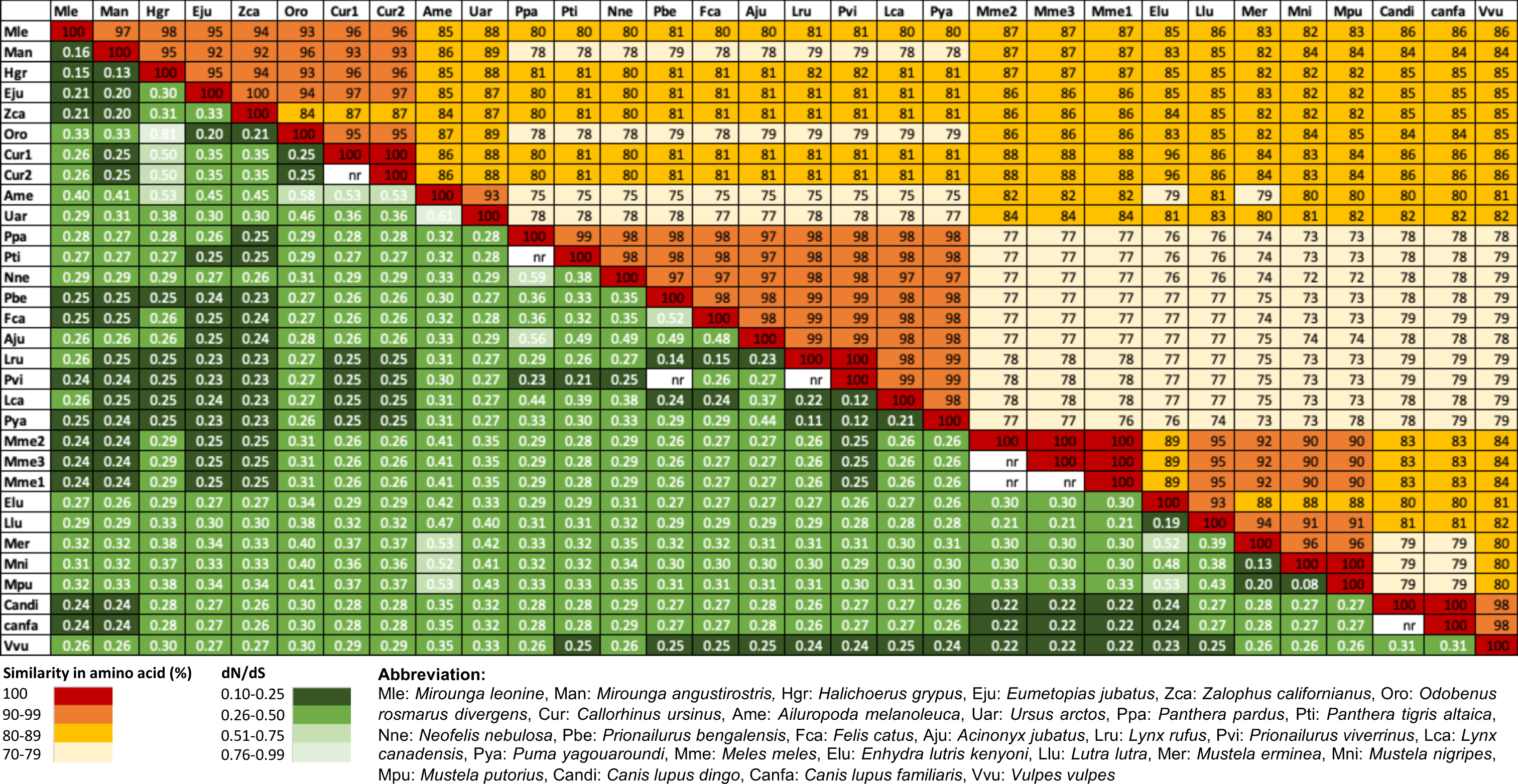
Sequence conservation and evidence for purifying selection of EnvV2 in carnivora. The table shows the pairwise percentages of amino acid sequence identity between EnvV2 and the indicated species in the upper triangular area, along with the non-synonymous/synonymous (dN/dS) rate. The pairwise Nei-Gojobori method (38) was used to construct the table. The nr code is due to the presence of only one or fewer non-synonymous and synonymous mutations between species; n.r. is not relevant. Color codes are provided below the table for both series of values.

As the newly identified EnvV2 in these species did not provide a complete long terminal repeat (LTR) sequence, we were unable to completely investigate the integration time into the host genome. Therefore, we constructed a TimeTree of mammalian vertebrates, as modified from a previous report (21, 44), with additional data on newly identified EnvV2 from other species. Based on the TimeTree analysis in Figure 7, we estimated that the integration time for newly identified EnvV2 during mammalian evolution, particularly in rodents, elephants, and bats, was similar to that of EnvV2 in Carnivora and ungulates; at least 60 million years ago (21). However, due to limited information on non-mammalian vertebrates, such as birds, reptiles, fishes, and amphibian, we were unable to confirm an integration time of EnvV2 in non-mammalian vertebrates in this study.

**Figure 7.**
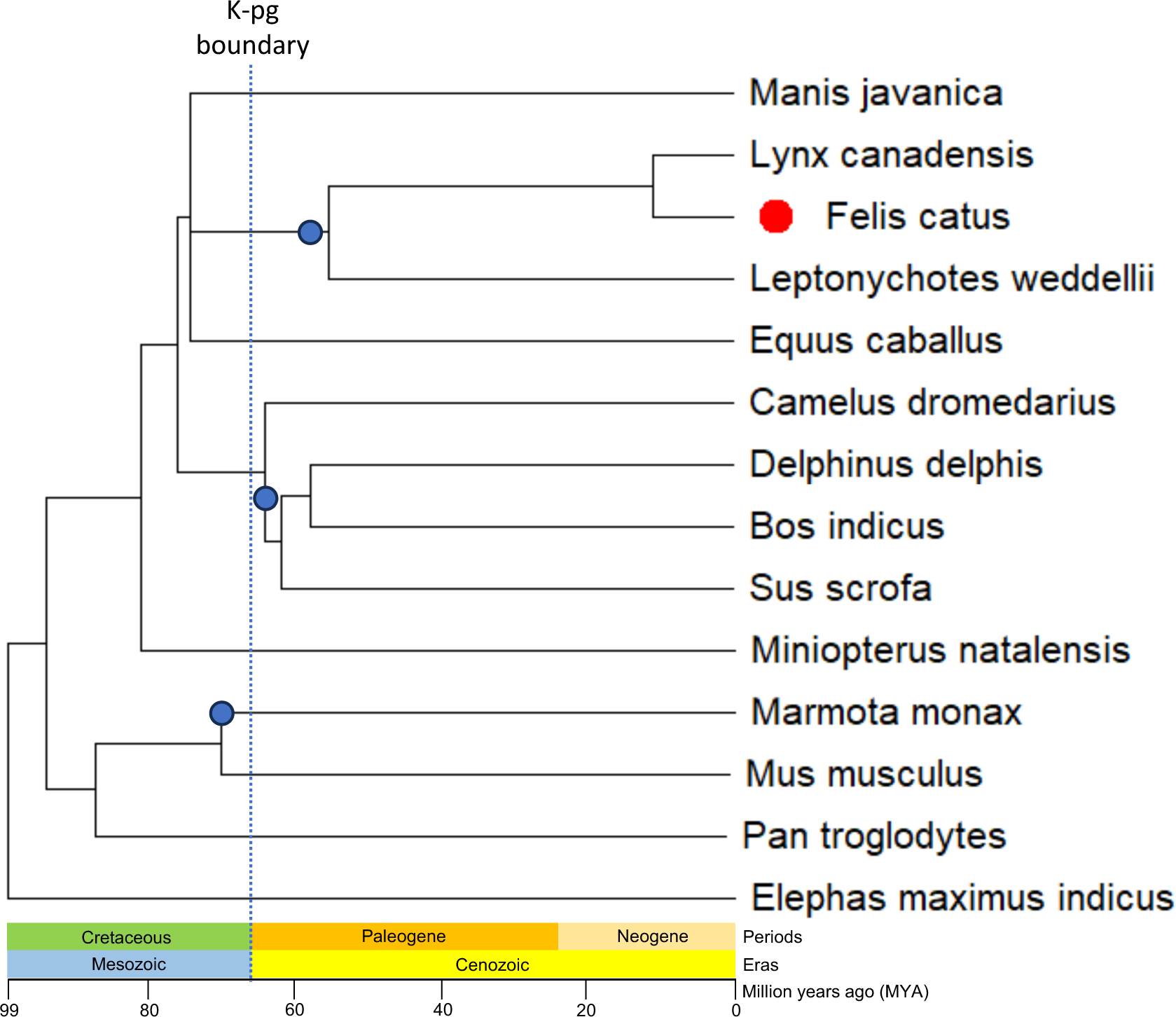
Evolution of ancient EnvV2 around the K-Pg period in mammalian vertebrates. (A) The cladogram illustrates the phylogenetic relationships among the indicated species belonging to mammalian vertebrates. The trees and divergence times were generated using TimeTree (44). Blue dots indicate possible insertion of EnvV2 into the host genome. The bar scale indicates million years ago (MYA).

## Discussion

In this study, while investigating the expression of ERV *env* genes in domestic cats, we discovered that EnvV2-Fca is closely related to the EnvV2-Hum gene (19). EnvV2-Fca comprises 439 amino acids, including a signal peptide, SU subunit, fusion peptide, and an ISD; however, the TM subunit is partially deleted (Figure 1A). In contrast, human and macaque monkey EnvV2 genes exhibit no structural defects. Expression analysis in culture cells indicated that EnvV2-Fca is released from cells as a soluble protein (Figure 2A), while the human and macaque monkey EnvV2 proteins are membrane bound (20). This solubility may be due to the partial deletion of the TM subunit in the Env protein, which is not anchored to the cell membrane and can be efficiently released from cells (14).

Expression analysis revealed that EnvV2-Fca is present in various feline tissues and cell lines (Figure 2B and 2C). The expression of EnvV2-Fca was observed at a similar level in tumor cell lines, specifically lymphoma and mammary carcinoma; however, cell type-specific expression was not observed in the cell lines used in this study. EnvV2-Fca expression in normal cells is currently unknown. In feline tissues, the highest level of EnvV2-Fca expression was detected in the placenta, as confirmed by qRT-PCR and in situ hybridization, and was specifically observed in the placental trophoblast syncytiotrophobic layer. These findings are similar to those of macaque monkey EnvV2-Mac, which is expressed in trophoblasts (20). This may be due to the evolutionary conservation of a regulatory mechanism of gene expression. When comparing EnvV2-Mac to EnvV2-Fca, we observed that EnvV2-Mac retained cell fusion activity (20), whereas EnvV2-Fca did not (Figure 4). The structural deletion in the TM subunit of the gene, resulting in the secretion of the EnvV2-Fca protein, may be the reason for the lack of fusion activity. However, EnvV2-Hum also lacks fusion activity (45), and considering that EnvV2-Hum exhibits no structural defects compared to EnvV2-Fca and Env-Mac (Figure 1), it is possible that the fusion ability of EnvV2-Fca was lost due to a genetic mutation, although the exact cause is unknown.

Our findings indicated that EnvV2 is widely present in many mammals (humans, primates, carnivora, even-toed ungulates, rodent, elephants, and bats), and a limited number of non-mammals (birds, reptiles, fishes, and amphibian) (Figure 5). As EnvV2 did not possess a complete LTR in these species, the integration time into the host genome, as determined using the LTR-based method (46), could not be evaluated. Based on the TimeTree analysis in Figure 7, the integration time for additional EnvV2 during mammalian evolution in rodents, elephants, and bats was estimated to be at least 60 million years ago, around the Cretaceous-Paleogene (K-Pg) boundary, which is similar to a previous study’s findings in carnivores and even-toed ungulates (21). However, we were unable to identify an integration time for non-mammalian vertebrates in this study due to limited information.

EnvV2 identified in mammals and non-mammals can be classified into two groups by genetic analysis: one contains a complete EnvV2, structurally similar to EnvV2-Hum and EnvV2-Mac, and the other is a defective EnvV2, similar to EnvV2-Fca (Figure 5). These structural features are consistent in each species. The partial deletion of the TM subunit, such as that observed in EnvV2-Fca, is commonly conserved in mammals (carnivores and even-toed ungulates) and is limited in non-mammals (birds, reptiles, fishes, and amphibian). Interestingly, some species (*Sciurus carolinensis*, *Aquila chrysaetos chrysaetos,* and *Gopherus evgoodei*) have two forms of EnvV2. However, it is difficult to determine the presence or absence of fusion activity based on structural features, as in EnvV2-Hum, EnvV2-Mac, and EnvV2-Fca. Notably, all identified EnvV2 proteins contained the ISD, except for that in *Rattus norvegicu*s. The ISD may have a suppressive effect on the immune response in cellular and humoral responses (47). Therefore, we speculate that EnvV2 may be involved in host immune tolerance through the maternal immune system to protect the fetus during pregnancy.

Currently, we have no evidence suggesting that EnvV2 has an inhibitory function against cell-fusion, similar to suppressyn’s effect on syncytin (16). EnvV2-Fca is specifically expressed in the placenta, suggesting that it may contribute to placentation via the regulation of cell-fusion activity. Comparing the structure of the defective EnvV2 group with Refrex-1 and suppressyn (14, 15), we can assume that the defected EnvV2 group may function as an antiviral factor against retroviral infections. Alternatively, the role of EnvV2 may be altered and play a different role in each species.

Studies have reported the potential cross-species transmission of ERVs (21, 48). The identification of EnvV2 in birds, bats, and rodents indicates the potential for the transmission of EnvV2 through these intermediate vectors. Bats are known to harbor several retroviruses, such as *Hervey pteropid gammaretrovirus*, which is closely related to koala retrovirus and Gibbon Ape Leukemia Virus (49). Bats have unique ecological characteristics, such as flight, hibernation, a relatively long lifespan, and colonial roosting behavior, which make them ideal vectors for the maintenance of viral infections (50). Rodents are also considered to be intermediate vectors for the spread of retroviruses (51, 52). Furthermore, birds are suspected to be involved in the transmission of retroviruses. Birds have been found to carry several retroviruses, including *gammaretrovirus*, reticuloendotheliosis virus (53), which are reported to be directly derived from mammalian retroviruses (54).

In conclusion, we found that EnvV2 evolution is not limited to mammals but extends to several non-mammalian vertebrates. Furthermore, EnvV2-Fca is a secreted protein and does not exhibit fusion activity. This suggests that EnvV2-Fca is not directly involved in syncytiotrophoblast formation during pregnancy. Instead, Syncytin-Car1 may be the primary mediator of this biological process in domestic cats (42). Further studies are needed to clarify the receptor and specific roles of the EnvV2-Fca protein. Notably, EnvV2 may have possible implications in immunotolerance during pregnancy or may act as a restrictive factor for exogenous retrovirus infection. Moreover, this study presents a scenario of retroviral transmission among species, which may have significant implications in our understanding of retroviral transmission and ERV evolution.

### Ethical approval

All animal experiments were conducted in accordance with the Guidelines for the Care and Use of Laboratory Animals, provided by the Ministry of Education, Culture, Sports, Science and Technology, Japan. All experiments were approved by the Genetic Modification Safety Committee of Yamaguchi University.

### Accession numbers

The accession numbers obtained from publicly available data are as follows: Human (NM_001191055), Macaque (NM_001348393), Chimpanzee (NM_001307974), *Felis catus* (XM_019831672), *Ursus arctos* (XM_057304973), *Panthera pardus* (XM_019425134), *Miniopterus natalensis* (XM_016197129), *Myotis daubentonii* (XM_059699531), *Myotis lucifugus* (XM_023753138), *Phyllostomus discolor* (XM_036014176), *Pipistrellus pipistrellus* (LR862361), *Anas platyrhynchos* (XM_038171956), *Aquila chrysaetos chrysaetos* (XM_041121140, XM_030010836), *Cygnus atratus* (XM_050710036), *Grus americana* (XM_054826583), *Indicator indicator* (XM_054389296), *Opisthocomus hoazin* (XM_009944966), *Castor canadensis* (XM_020155210), *Ictidomys tridecemlineatus* (XM_040273522), *Marmota monax* (XM_046457090), *Rattus norvegicus* (NM_001408907), *Sciurus carolinensis* (XM_047551960, XM_047551962), *Bos indicus* (XM_019957350), *Bubalus bubalis* (XM_025288582), *Alligator mississippiensis* (XM_059724950), *Gopherus evgoodei* (XM_030542918, XM_030542367), Oncorhynchus mykiss (XM_036942505), *Syngnathus scovelli* (XM_049756540), *Elephas maximus indicus* (XM_049877704), *Rana temporaria* (XM_040342020). The sequences from the isolate described in this paper have been deposited in DDBJ/EMBL/GenBank under accession number LC794385.

## Acknowledgments

We are grateful to Dr. Takayuki Nakagawa (The University of Tokyo) for providing feline mammary adenocarcinoma cells (FMC, FKNp, FON, FRM, and FNNm). We would like to thank Editage (www.editage.jp) for English language editing.

## Author Contributions

The contributions of the authors are described as follows. Conceptualization: K.N., Data Curation: D.P.,K.S.,A.M.,K.N., Formal analysis: D.P.,K.S.,K.N., Funding acquisition: K.N., Investigation: D.P.,K.S.,A.M.,K.N., Methodology: D.P.,K.S.,A.M.,K.N., Project administration: K.N., Supervision: T.K.,K.N., Validation: D.P.,K.S.,A.M., K.N., Writing-original draft: D.P.,K.N., Writing-review & draft: D.P.,K.N.

